# Distinct human stem cell subpopulations drive adipogenesis and fibrosis in musculoskeletal injury

**DOI:** 10.1101/2023.07.28.551038

**Authors:** Steven M. Garcia, Justin Lau, Agustin Diaz, Hannah Chi, Miguel Lizarraga, Aboubacar Wague, Cristhian Montenegro, Michael R. Davies, Xuhui Liu, Brian T. Feeley

## Abstract

Fibroadipogenic progenitors (FAPs) maintain healthy skeletal muscle in homeostasis but drive muscle degeneration in chronic injuries by promoting adipogenesis and fibrosis. To uncover how these stem cells switch from a pro-regenerative to pro-degenerative role we perform single-cell mRNA sequencing of human FAPs from healthy and injured human muscles across a spectrum of injury, focusing on rotator cuff tears. We identify multiple subpopulations with progenitor, adipogenic, or fibrogenic gene signatures. We utilize full spectrum flow cytometry to identify distinct FAP subpopulations based on highly multiplexed protein expression. Injury severity increases adipogenic commitment of FAP subpopulations and is driven by the downregulation of DLK1. Treatment of FAPs both *in vitro* and *in vivo* with DLK1 reduces adipogenesis and fatty infiltration, suggesting that during injury, reduced DLK1 within a subpopulation of FAPs may drive degeneration. This work highlights how stem cells perform varied functions depending on tissue context, by dynamically regulating subpopulation fate commitment, which can be targeted improve patient outcomes after injury.

## Introduction

Musculoskeletal injuries impact millions of people each year, are an enormous burden on the health care system, and cause significant morbidity in these patient populations^1^. In this study, we sought to investigate stem cell functional heterogeneity in human rotator cuff injury, a clinically relevant model of human muscle disease that is also directly applicable to many other muscle degenerative pathologies. Rotator cuff injuries are the most common chronic upper extremity problem in the United States and affect more than 4.5 million people per year^1,2^. In rotator cuff tears, poor clinical outcomes and high retear rates are directly correlated with increased fat and fibrosis and muscle atrophy, termed fibro/fatty degeneration^3^. Recent studies of rotator cuff muscle degeneration suggest this degeneration is mediated by fibroadipogenic progenitor cells (FAPs)^4^. Understanding the molecular and cellular drivers of fibro/fatty degeneration and muscle atrophy is critical for improving muscle regeneration after rotator cuff injury and repair. The mechanisms underlying fibro/fatty degeneration in rotator cuff injury may also be translatable to other muscle degenerative states including aging and muscular dystrophy^5,6^.

A comprehensive understanding of human FAP cell heterogeneity and potential functional FAP subpopulations is limited by the challenges of obtaining normal and injured human tissue. Several studies of human skeletal muscle have shown unique FAP populations, but these studies have been limited by cell and patient number^7–9^. Furthermore, studies in a clinically relevant chronic injury model are lacking. In healthy tissue, FAPs, identified by expression of PDGFRα, regulate and promote myogenic regeneration^10,11^. In chronic injury, however, FAPs impede regeneration by differentiating into adipocytes and fibroblasts and secreting factors that promote adipogenesis and fibrosis, resulting in fibro/fatty degeneration^9–12^. Single cell RNA sequencing (scRNA-seq) studies in both mouse and human tissues provide evidence that subpopulations of PDGFRα+ cells exist in adult tissues, including skeletal muscle^7–9,12–16^. A recent investigation of adult mouse skeletal muscle found multiple unique subpopulations, several of which seem to be responsible for *de novo* adipogenesis and tissue mineralization^16^. Heterogeneous FAP subpopulations may therefore explain the multitude of roles these cells play in regenerating tissue; however, it is unclear if this holds true in humans.

In this study, we performed scRNA-seq of over 24,000 human FAPs from healthy tissue and injured rotator cuff muscles across a spectrum of injury. We generated a robust dataset of 14 unique muscle samples, including 6 patients with paired injured rotator cuff muscle and healthy deltoid muscle samples taken together at the time of surgery. We identified FAP heterogeneity through gene and protein expression and discovered how specific FAP subpopulations undergo transcriptional changes that correlate with injury severity and promote production of degenerative adipogenic tissue during injury. Utilizing high-dimensional full spectrum flow cytometry we identify distinct progenitor, adipogenic, and fibrogenic FAP subpopulations by multiplexed protein expression. We uncover that injury severity increases adipogenic commitment of FAP subpopulations and that this is driven by downregulation of Delta Like Non-Canonical Notch Ligand 1 (DLK1). Treatment of FAPs with DLK1 reduced adipogenesis *in vitro*, suggesting that during injury, reduced DLK1 within a subpopulation of FAPs may drive adipogenic degeneration. We then demonstrate in a xenotransplant model of human FAPs that treatment with DLK1 after injury results in less fatty infiltration *in vivo*. These experiments highlight the significance of stem cell heterogeneity in clinically relevant human samples. Our findings that tissue degeneration is driven by the regulation of specific stem cell populations within the larger stem cell pool presents opportunities for development of targeted therapies to improve patient outcomes for musculoskeletal injuries.

## Results

### Single cell RNA-sequencing identifies distinct human fibroadipogenic progenitor populations

We evaluated FAP heterogeneity in muscle biopsies from patients undergoing rotator cuff repair. We obtained biopsies from injured supraspinatus muscle with either a partial or full thickness tear and from non-injured deltoid muscle taken from the same patients, as healthy control tissue. As an additional source of healthy muscle, we obtained hamstring (semitendinosus and gracilis) muscle biopsies (**Figure 1a**). Samples were enriched for human FAPs by the surface marker combination CD45-/CD31-/CD56-/CD34+/PDGFRα and single cell RNA sequencing was performed (**Figures 1a and S1a**). CD34 has previously been shown to mark PDGFRα human FAPs^9^. We integrated all single cell datasets and identified a total of 7 major cell types by enriched expression of known marker genes including: endothelial cells (*CD31*), pericytes (*RGS5*), fibroblast/myofibroblasts (*ACTA2*), myogenic cells (*PAX7 & MYOD1*), T-cells (*CD3E*), macrophages (*CD68*), and FAPs (*PDGFR*α) (**Figures 1b, S1b, S1c and Supplementary Table S1**). The majority of cells captured (24,929 out of 39,356) expressed high levels of *PDGFRα*. Therefore, we robustly enriched for and profiled human FAPs from injured and non-injured muscle.

**Figure 1:**
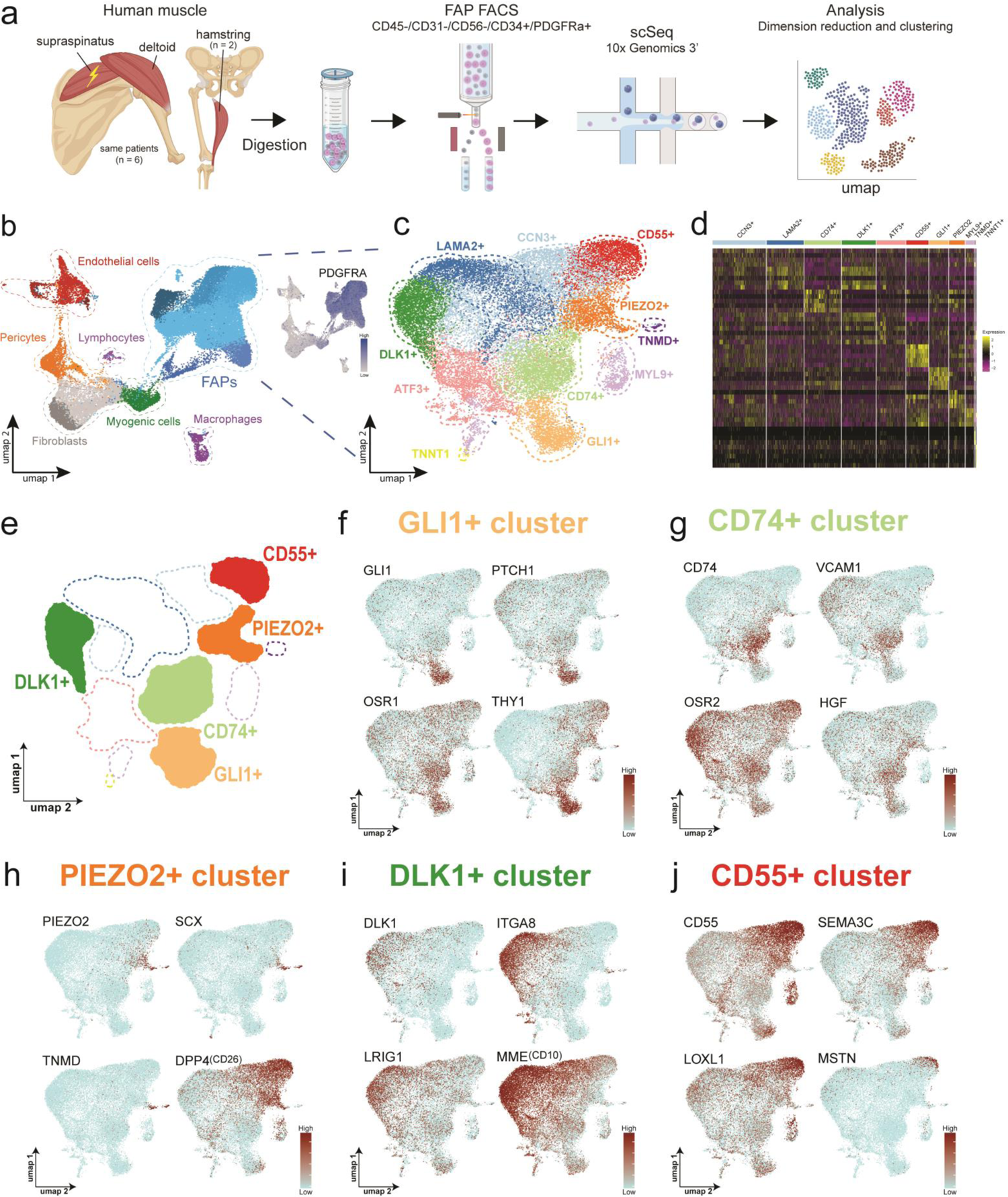
Single cell sequencing identifies distinct human fibroadipogenic progenitor populations. **a)** Dissociation, isolation, and single cell mRNA sequencing (scRNA-seq) of *PDGFRα*-expressing fibroadipogenic progenitors from human supraspinatus, deltoid, and hamstring muscle. **b)** UMAP of all cells (39,356). Color indicates cell type. Inset: *PDGFRα* expression. Color indicates expression level. **c)** UMAP of PDGFRα+ FAPs (24,929 cells). Color and dashed lines show cluster identities. Clusters are named by a top enriched gene markers for that population. **d)** Heatmap of top enriched genes from each FAP subpopulation. Colored bar indicates cluster identity. **e)** Model UMAP of FAP subpopulation with highlighted populations. **f-j** Expression plots of upregulated genes in the **f)** GLI1+, **g)** CD74+, **h)** DLK1+, **i)** PIEZO2+, and **j)** CD55+ populations. Deeper red indicates higher expression, scaled per plot. See also Supplementary Figure 1 and 2.

We next subclustered the *PDGFRα*+ FAPs and identified 11 transcriptomically unique FAP subpopulations (**Figures 1c, 1d, S1d and Supplementary Table S2**). The FAP subpopulations were named after a top differentially expressed marker in each respective population (**Figures S1e and S1f**). A GLI1+ cluster was notable for high and specific expression of hedgehog signaling genes (*GLI1* and *PTCH1*) and markers of stem/progenitor FAPs (*THY1* and *OSR1*), suggesting that cells in this cluster function as a FAP stem/progenitor cells (**Figures 1e and 1f**)^16–18^. A CD74+ population expressed markers of activated progenitor FAPs, including *VCAM1*, *OSR1* and *OSR2*, and *HGF* (**Figure 1g**). VCAM1 marks activated FAPs in adult mice, while OSR1 and OSR2 mark embryologic FAP progenitor populations^13,19^. HGF is a ligand involved in the expansion of progenitor populations during muscle regeneration^20,21^. These findings suggest that the CD74+ population functions as activated progenitors.

We identified a cluster expressing *PIEZO2*, a receptor involved in mechanosensation, suggesting that a subpopulation of FAPs may have mechanosensory function akin to other skeletal muscle stem cell populations (**Figure 1h**)^22–24^. This population also expressed markers of tendon progenitor cells including *SCX, TMND*, and *DPP4 (CD26)*^25^. The PIEZO2+ population may serve as a mechanosensitive population that gives rise to tendon progenitor cells. PIEZO2+ FAPs could be relevant in rotator cuff pathology, given the mechanical properties of rotator cuff muscles are altered with increasing severity of tear.

We also found two FAP clusters marked by *DLK1* or *CD55*, with upregulation of genes involved in either adipogenesis or fibrosis respectively. DLK1 is a known marker of adipose progenitors and has been studied in adipose tissue (**Figure 1i**)^26–28^. MME(CD10), a marker of human adipogenic FAPs^29^, and *ITGA8* and *LRIG1,* genes that increase during adipogenic commitment and differentiation, were all highly expressed in DLK1+ cells^30,31^. The CD55+ population was marked by expression of DPP4 (CD26) (**Figure 1h**); as well as, pro-fibrotic genes, such as *SEMA3C*, *LOXL1*, and *CD248* (**Figure 1j and Supplementary Table S2**)^32–34^. CD55+ cells also express *MSTN* (myostatin), a driver of muscle atrophy, suggesting this population of FAPs may be involved in the regulation of muscle growth and development^35^. Taken together, we hypothesize that DLK1+ and CD55+ cells represent subpopulations of FAPs that are primed to become either adipocytes or fibroblasts, respectively.

Additionally, we identified a TNMD+ myotendinous progenitor population, two myofibroblast clusters marked by MYL9+ and TNNT1+, and two transitional clusters marked by CCN3+ and LAMA2+ (**Figures S2a and S2b)**.

This dataset demonstrates the heterogeneity of human FAPs and highlights two main FAP subpopulations that express markers associated with adipogenesis or fibrosis. Expression of adipogenic progenitor genes in the DLK1+ population and pro-fibrotic genes in the CD55+ population, suggests that the dual function of FAPs as pro-fibrotic or pro-adipogenesis may derive from discrete subpopulations of FAPs poised to give rise to either fibroblasts or adipocytes^26,29–33^.

### Transcriptionally heterogenous human FAPs also have heterogenous protein expression of distinct surface markers

Our scRNA-seq human FAP dataset revealed specific expression of genes that encode surface proteins in FAP subpopulations. Co-staining PDGFRα with the surface markers THY1, DLK1, and CD55 confirmed human FAPs have heterogeneous expression of the surface marker proteins identified as population markers in the scRNA-seq analyses (**Figure 2a**). We hypothesized we could label and isolate FAP subpopulations based on the expression of these markers with flow cytometry. To rigorously test whether the transcriptionally-defined FAP subpopulations could be identified at the protein level, we utilized full spectrum flow cytometry. In contrast to conventional flow cytometry, which is limited to a handful of fluorescent markers, full spectrum flow cytometry permits high dimensional profiling of up to 40 fluorescence markers^36^. We used this technique to profile the expression of surface markers in FAPs derived from healthy human muscle (**Figure 2b**). Live (SYTOX-) FAPs were identified by the marker combination CD45-/CD31-/CD56-/CD34+ and further profiled with 6 additional surface markers that were enriched in four different FAP subpopulations. CD74 (CD74+ cluster), DLK1 and CD10 (MME) (DLK1+ cluster), CD55 and CD26 (DPP4) (CD55+ cluster), and THY1 (GLI1+ cluster)(**Figures S3a, S3b, and 2c**).

**Figure 2:**
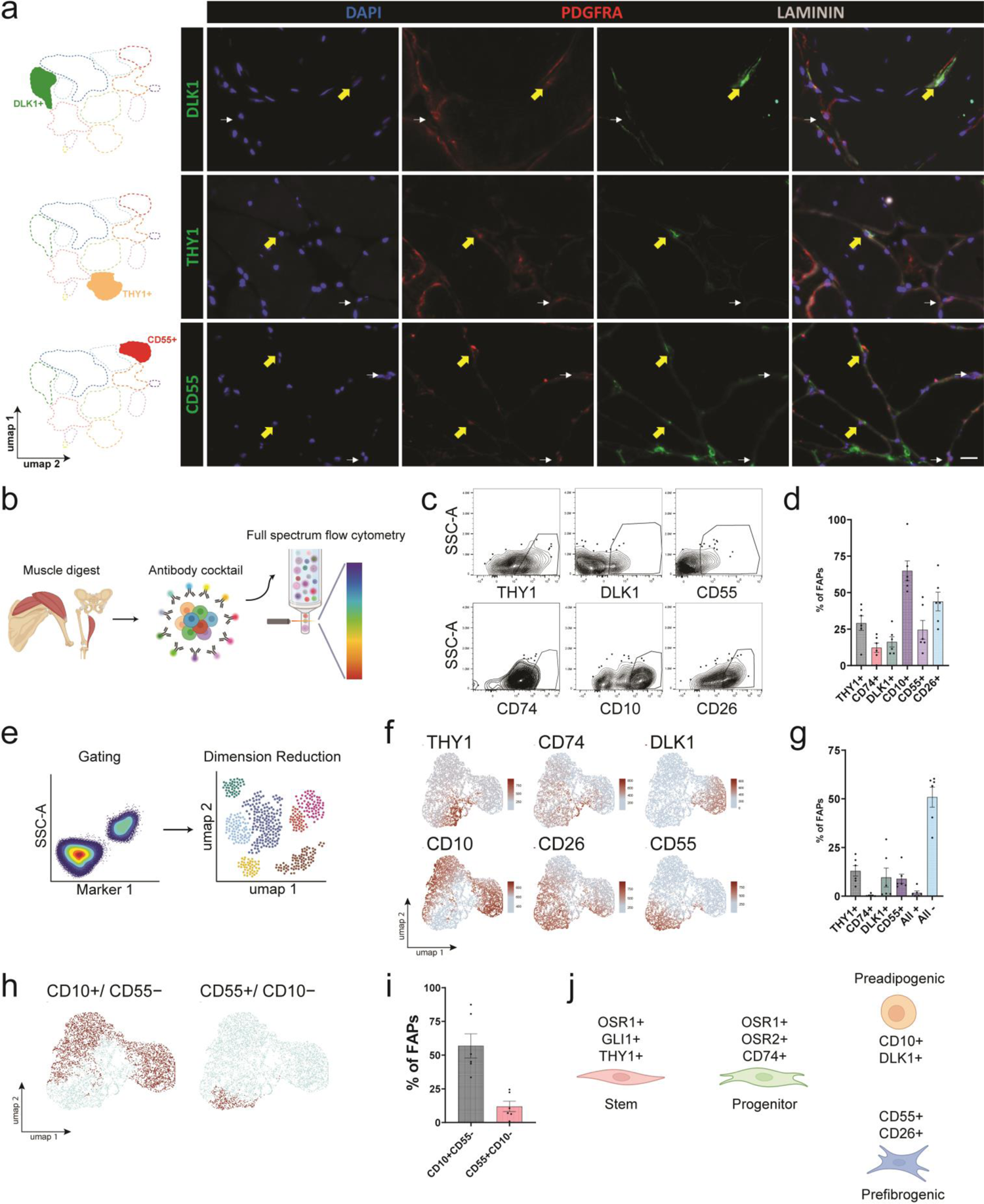
Heterogenous expression of distinct surface marker proteins in human fibroadipogenic progenitors. **a)** Left: Model UMAP plots highlighting population of interest. Right: multiplexed immunofluorescence for DLK1 (top panels), THY1 (middle panels), and CD55 (bottom panels) with PDGFRα on human muscle cross sections. Large yellow arrows: PDGFRα+ cells positive for the population marker. Small white arrows: PDGFRα+ cells negative for the population marker. Scale bar = 20 µm. n = 3 biological replicates. **b)** Experimental design of full spectrum flow cytometry analysis of human FAPs. The antibody cocktail included antibodies against CD31, CD45, CD56, CD34, PDGFRα, THY1, CD74, DLK1, CD10, CD55, CD26, and a SYTOX live dead stain. **c)** Representative full spectrum flow cytometry plots showing FAP surface protein expression. **d)** Percentage of FAPs expressing each surface protein. Percentage is calculated by expression of each protein in all FAPs, independent of expression of other proteins. **e)** Schematic of dimensional reduction of full spectrum flow cytometry data. Expression of all multiplexed surface proteins were input in the UMAP algorithm. **f)** Representative UMAPs of full spectrum flow cytometry data. Color indicates expression of protein marker. **g)** Percentage of FAPs with exclusive expression of each protein, expression of all proteins, or lack of expression of all proteins. **h)** UMAP with CD10+ and CD55+ cells highlighted in red. Each cell was binned into either positive or negative for each marker based on expression thresholds for CD10 and CD55. **i)** Percentage of FAPs that are CD10+CD55- and CD10-CD55+. **j)** Model of human FAP subpopulations with the surface marker profiles of each population. n = 6 biological replicates for all data in b-i. All plots represent mean ± SEM. See also Supplementary Figure 3.

To first confirm expression of each protein, we determined the percentage of FAPs expressing each surface protein (**Figure 2d**). We then performed dimensional reduction on the expression levels of all 6 proteins (**Figure 2e**)^37^. Clustering analysis based on the expression of these 6 surface proteins resulted in similar clusters to those observed in the scRNA-seq analysis (**Figure 2f**). THY1+ cells clustered together and partially overlapped CD74+ cells. We identified CD55 expressing cells and a subset of CD55+ cells that co-expressed CD26, which matched the *CD55* and *CD26* transcript expression patterns (**Figure S3c**). Additionally, we found two populations of CD10+ cells, one that did not express DLK1 and a second population that co-expressed DLK1, again in concordance with our scRNA-seq data (**Figures S3d, S3e, and S3f**). We quantified the percentage of FAPs belonging to each of the major FAP subtypes: THY1+, DLK1+, and CD55+ FAPs (**Figures 2g and S3f**). Importantly, this data demonstrates that protein expression of CD55 and CD10 are largely mutually exclusive, suggesting that these clusters represent FAPs in distinct cell states (**Figure 2h and 2i**). We hypothesize expression of THY1+, DLK1+/CD10+, or CD55+ promotes FAPs to adopt progenitor, pre-adipogenic, or pre-fibrogenic cellular fates, respectively (**Figure 2j**). Overall, this data confirms that FAP transcriptional heterogeneity is translated into distinct protein expression patterns.

### DLK1/CD10 and CD55 are upregulated during FAP differentiation towards adipocyte or fibroblast cell fates

Previous work has suggested that a subpopulation of FAPs expressing *OSR1* function as progenitors for both fibroblasts and adipocytes^16,17,19,38^. We therefore tested our hypothesis that FAP subpopulations include FAP progenitors (marked by *GLI1, THY1*, and *OSR1*) and activated progenitors (marked by *CD74*, *OSR1*, and *OSR2*) which differentiate to FAPs committed to the fibroblast lineage (marked by *CD55*), adipocyte lineage (marked by *DLK1* and *CD10*), or tenogenic lineage (marked by *TNMD*). Pseudotime analysis revealed a path from GLI1+ FAPs to *CD55*, *DLK1*, or *TNMD* expressing FAPs (**Figures 3a and S4a**). We found increasing expression of surface markers for FAP subpopulations (*DLK1*, *CD55*, and *CD26*) and genes associated with adipogenic, fibrogenic, or tenogenic identity (**Figures 3b and S4b**). We then focused on the lineages leading from FAP progenitors to pre-adipogenic and pre-fibrogenic FAPs, as increased fibrosis and adipogenesis characterizes typical muscle degeneration.

**Figure 3:**
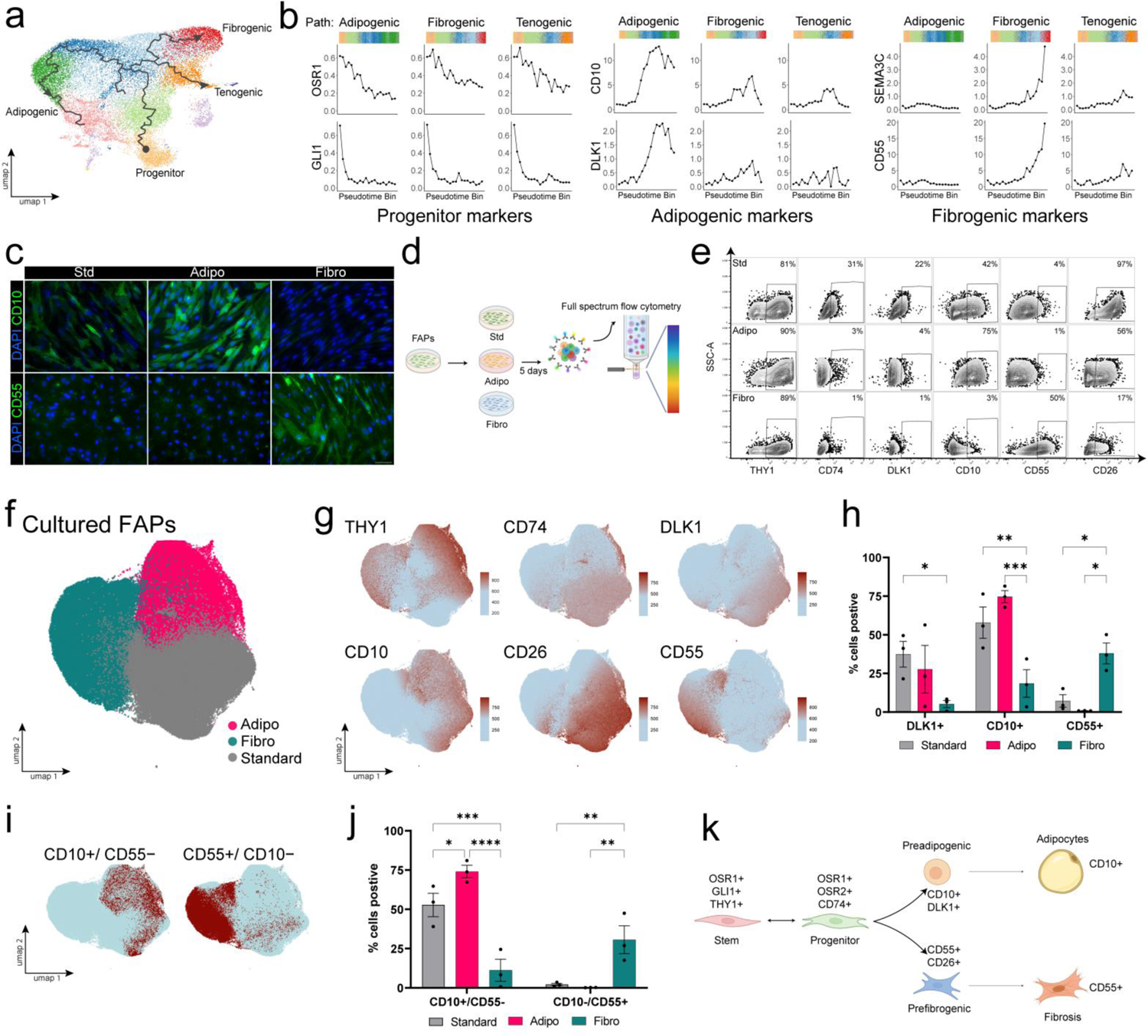
DLK1/CD10 and CD55 are upregulated during FAP differentiation towards adipocyte or fibroblast cell fates. **a)** UMAP of all FAPs with pseudotime trajectories from progenitor cells to adipogenic, fibrogenic, and tenogenic FAPs. Color indicates cluster identity. **b)** Binned average expression of select genes along each pseudotime trajectory leading to an adipogenic, fibrogenic, or tenogenic fate. Colored bars indicate the cluster identity for each cell along the trajectory. **c)** Immunofluorescence images of FAPs grown for five days in either standard expansion (std), adipogenic (adipo), or fibrogenic media (fibro). Scale bar = 20 µm. **d)** Experimental design of human FAP full spectrum analysis of cultured cells. **e)** Representative flow cytometry plots of FAP surface protein expression. **f)** UMAP of all cultured cells. Color indicates media condition. **g)** Surface protein expression overlaid on UMAP. Color indicates level of protein expression. **h)** Percentage of FAPs expressing DLK1, CD10, or CD55 in each media condition. **i)** UMAP with CD10+ and CD55+ cells highlighted in red. Each cell was binned into either positive or negative for each marker based on expression thresholds for CD10 and CD55. **j)** Percentage of CD10+CD55- and CD10-CD55+ FAPs in each media condition. **k)** Model of proposed human FAP lineages and expression of select genes. n = 6 biological replicates for all data. All plots represent mean ± SEM. Statistical test for all plots: two-way ANOVA with Tukey’s multiple comparison correction. *p < 0.05 **p < 0.01 ***p < 0.001. See also Supplementary Figure 4.

We first asked how FAP subpopulations are transcriptomically altered when directed towards a fibrogenic or adipogenic fate. We performed scRNA-seq of FAPs cultured for 3 weeks in either adipogenic, fibrogenic, or a standard base expansion media that maintains FAPs in a proliferative state (adipogenic condition reanalyzed from Davies et al.^39^) (**Figures S4c**). Cells from each condition largely grouped together (**Figures S4c and Supplementary Table S3**). Expression of *PDGFRα*, *PLIN1*, and *ACTA2*, was expressed predominantly in cells cultured in the standard base media, adipogenic media, and fibrogenic media, respectively (**Figure S4d**).

We found FAPs maintained in standard base media expressed the stem/progenitor markers including *OSR1* and *CD74* (**Figure S4e and Supplementary Table S4**). FAPs cultured in adipogenic media produced a group of adipocytes, marked by expression of the adipocyte marker *PLIN1* and absence of FAP marker *PDGFRα* (**Figure S4e**). Similarly, FAPs cultured in fibrogenic media produced a group of fibroblasts, marked by *ACTA2* and decreased *PDFGRα* (**Figure S4e**). FAPs cultured in adipogenic media expressed higher levels of *CD10*, while FAPS cultured in fibrogenic media expressed higher levels of CD55, supporting the hypothesis that FAPs marked by *CD10* and *CD55* are pre-adipogenic and pre-fibrotic FAPs (**Figure S4f**). Thus, human FAPs could be directed through an intermediate pre-adipogenic or pre-fibrotic FAP state towards *PLIN1*+ adipocyte or *ACTA2*+ fibroblast fate.

We next asked if we could capture FAPs as they transition from progenitors to either adipocytes or fibroblasts and identify their cellular state by protein surface expression. We found FAPs increase protein expression of CD10 in adipogenic media and CD55 in fibrogenic media, respectively, after 5 days in standard, adipogenic, or fibrogenic media conditions (**Figure 3c**). We analyzed FAPs with full spectrum flow cytometry for our panel of FAP surface markers; CD74 (CD74+ cluster), DLK1 and CD10 (DLK1+ cluster), CD55 and CD26 (CD55+ cluster), and THY1 (GLI1+ cluster) (**Figures 3d and 3e**). We also measured surface protein expression in FAPs that were not cultured, termed freshly harvested FAPs. After dimensional reduction, FAPs largely clustered together by the media they were grown in, with some overlap, suggesting that the culture media produces distinct expression patterns of these markers (**Figure 3f**).

As in our scRNA-seq dataset of human FAPs in Figures 1 and 2, DLK1+ cells were identified as a subset of CD10+ cells (**Figure S4h**). The adipogenic marker CD10 was largely expressed by cells in the adipogenic treatment group, while the fibrogenic marker CD55 was largely expressed by cells in the fibrogenic group (**Figures 3h**). We found CD10+CD55-FAPs were near exclusive to adipogenic media conditions while conversely, CD10-CD55+ FAPs were exclusive to fibrogenic media (**Figure 3i and 3j**). Compared to freshly harvested FAPs from healthy muscle, these two cell subsets were all also increased in each media respectively (**Figure S4i**). Taken together, the specific expression of CD10 and CD55 suggests that these two surface proteins can been utilized to mark human pre-adipogenic and pre-fibrogenic FAPs. The combination of mRNA and protein data at single cell resolution suggests that two FAP subpopulations contribute to the production of fibrosis and fatty infiltration in human muscle. This data also identifies multiple markers of FAPs as they progress from the progenitor state to distinct lineages (**Figure 3k**).

### Severe chronic injury increases adipogenic and fibrotic commitment of FAPs

We next sought to assess FAP subpopulations in a clinically significant model of human muscle disease: rotator cuff injuries. Rotator cuff injuries occur on a spectrum from partial thickness (incomplete) to full thickness (complete) tendon tears (**Figure 4a**)^2^. This spectrum of tendon injury is correlated with the severity of muscle degeneration in chronic injuries. Chronic full thickness tears readily develop fatty degeneration, while partial tears do not^40^. The degree of degeneration can be quantified clinically with magnetic resonance imaging (MRI) (**Figure 4b**). As degeneration increases, the degree of adipogenic infiltration is classified on MRI from Goutallier 0 (no fat), Goutallier 1-2 (moderate fat), and Goutallier 3-4 (severe fatty degeneration)^41,42^. To investigate the differences in chronic muscle injury, we compared chronically injured rotator cuff muscle to control healthy deltoid muscle taken from the same patients. Patients with full thickness injuries had moderate muscle fatty infiltration while partial tear patients’ muscle quality were all without fatty infiltration (patients with severe, Goutallier 3-4, fatty degeneration generally are not candidates for repair) (**Figures 4b and S5a**).

**Figure 4:**
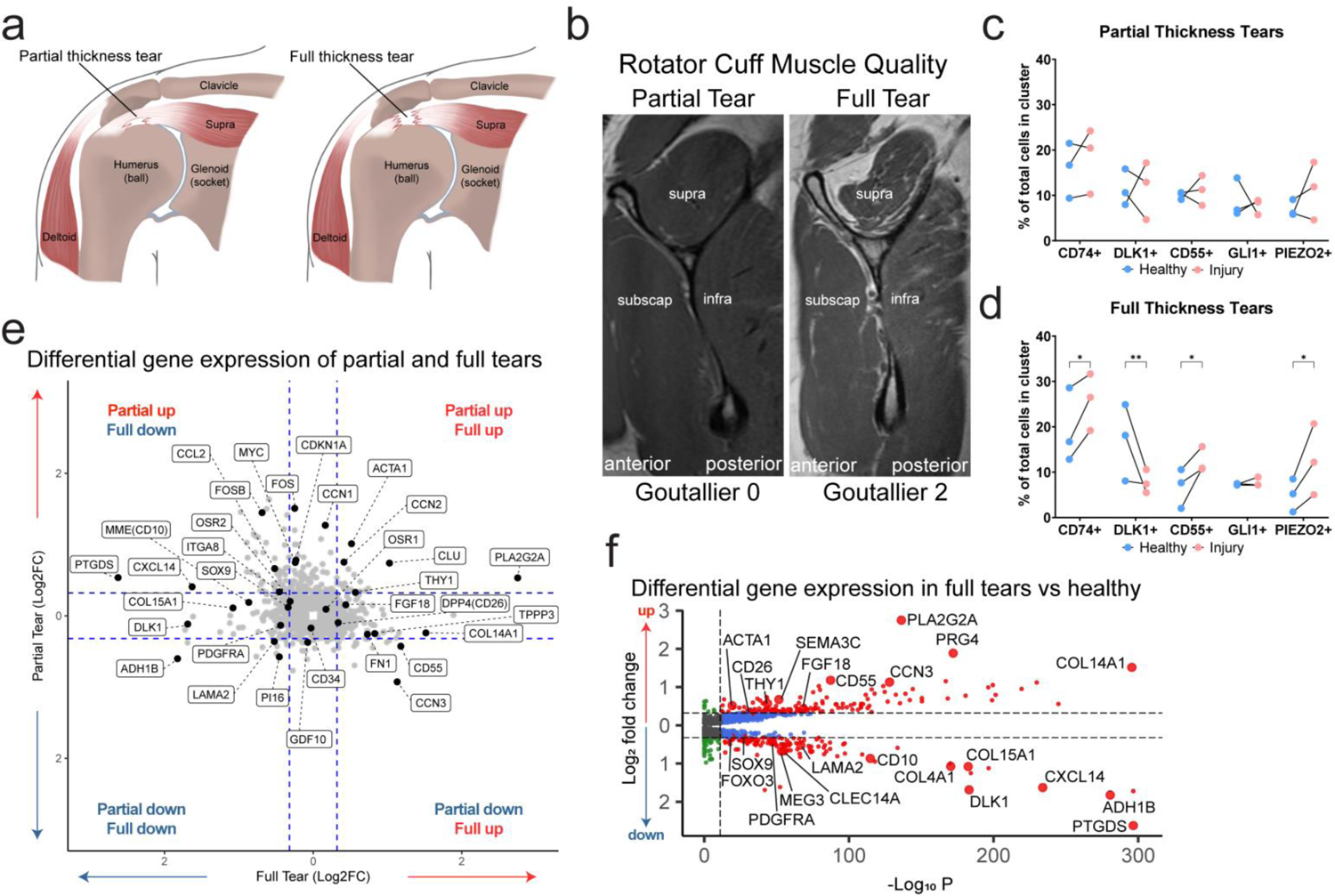
Severe chronic injury increases adipogenic and fibrotic commitment of FAPs. **a)** Model cross-section of the human shoulder depicting partial (incomplete thickness) or full (complete thickness) rotator cuff tendon tears. **b)** Representative cross-section, T2-weighted, MRI images of human rotator cuff muscles. Supra (supraspinatus), infra (infraspinatus), subscap (subscapularis) muscles. Fatty infiltration classification: Goutallier 0-4. **c-d)** The proportion of each cluster in paired healthy and injured muscle samples for: **c)** partial thickness tendon tears and **d)** full thickness tendon tears compare to healthy deltoid muscle from the same patients. Color indicates if cells are from healthy (blue) or injured (pink) muscle sample. Paired samples linked with lines. n = 3 partial and n = 3 full thickness tear biological replicates. Statistically significant tested with a paired two-way ANOVA. *p < 0.05 **p < 0.01. **e)** Scatter plot showing gene Log_2_FC in FAPs derived from muscles with full tears compared to FAPs derived from healthy muscles (x-axis) versus Log_2_FC in FAPs derived from muscles with partial tears compared to FAPs derived from healthy muscles (y-axis). Significant expression changes were Log_2_FC > 0.32 (fold change > 1.25) and p < 0.05 (blue dotted lines). **f)** Volcano plot demonstrating significant gene differential expression in full thickness tear muscle samples compared to healthy control muscle. Significant genes were Log_2_FC > 0.32 (fold change > 1.25) and p < 0.05 (red circles). See also Supplementary Figure 5.

We first compared the proportion of FAP subpopulations between healthy and injury groups in our FAP scRNA-seq dataset from Figure 1 (**Figure S5b**). The proportion of each FAP subpopulation did not change in patients with partial thickness tears in comparison to healthy controls (**Figures 4c and S5c**). However, patients with full thickness tears had altered proportions of FAP subpopulations compared to control. We found an increased proportion of activated CD74+, pre-fibrogenic CD55+, and mechanosensitive PIEZO2+ FAP subpopulations, and a decreased proportion of pre-adipogenic DLK1+ and transitional CCN3+ FAP subpopulations (**Figures 4d and S5d**). Thus, in contrast to partial thickness tears, full thickness tears result in altered FAP subpopulation composition. Full thickness tears resulted in increased proportions of activated progenitor, fibrogenic, and mechanosensitive cells. Interestingly, the increased proportion of PIEZO2+ cells suggest these cells could function as tendon progenitor cells^43^. Full thickness tendon injuries produce mechanical changes within the muscle, such as unloading, which may push FAPs toward this lineage. Together, this data suggests that changes in FAP subpopulation composition may drive the fibrotic and fatty degeneration specific to full thickness tears.

We next performed differential gene expression analysis of FAPs from the partial thickness and full thickness injured supraspinatus muscle groups compared to each patient group’s corresponding control deltoid muscles. Differentially expressed genes were compared between the two injuries (**Figure 4e and Supplementary Tables S5 and S6**). We found genes that were up- or down-regulated in both injuries (bottom left and top right quadrants), and genes that were altered differentially between the two injuries (top left and bottom right quadrants). Notably *THY1* and *PLA2G2A* expression were significantly upregulated in both partial and full tears. *THY1* has been shown to be upregulated in human FAPs from the muscle of diabetic patients with adipogenic degeneration and *PLA2G2A* expression is increased in obesity^9,44^. Cell activation related genes, *FOS*, *MYC*, and *CDKN1A*, were significantly upregulated in the partial tear group compared to the full tear group. Genes involved in pro-adipogenic and pro-myotendinous regeneration were upregulated as injury progressed from partial to full tear (**Figure 4e**). Genes associated with repression of adipogenesis, such as *DLK1* and *SOX9*, were significantly downregulated in the full thickness tear group but not in the partial tear group (**Figures 4e and 4f**), while genes associated with collagen remodeling and myotendon regeneration, such as *TPPP3*, *FN1*, *FBN1*, and *COL14A1*, were significantly upregulated in the full thickness tear group but not in the partial tear group (**Figures 4e and 4f**)^25,43^. These results correlate with the degree of fatty degeneration seen on MRI, where only full thickness tears show histologically increased adipogenesis and fibrosis^26,45,46^. These data suggest that FAP populations are altered on a continuum depending on injury severity and may explain the clinical changes in muscle function and structure seen in these patient groups.

### FAP subpopulation gene expression in chronic injury leads to a predominantly pro-adipogenic tissue state

We hypothesized that the altered proportions of FAP subpopulations and gene expression changes in injured muscle are reflective of altered FAP lineage commitment. To investigate this, we utilized RNA velocity to map cell trajectories in both healthy muscle and chronic full thickness tears (**Figures 5a, 5b, and S6a**)^47^. In healthy muscle, the major cell trajectories arise from the OSR1+/OSR2+/CD74+ progenitor population which splits into 2 pathways towards the pre-adipogenic (DLK1+) and pre-fibrogenic (CD55+) populations (**Figure 5a**). Cells emanating from the mechanosensitive (PIEZO2+) population are directed towards either the pre-fibrogenic or tenocyte (SCX+/TNMD+) progenitor population. We found FAPs from full thickness tear muscle had shifted cell trajectories towards the pre-adipogenic cells (**Figure 5b**). The pathway bifurcation from the CD74+ progenitor population, seen in healthy muscle FAPs, was reduced to a single pro-adipogenic pathway (**Figures 5b**). Taken together, this data suggests the cells in the progenitor subpopulation are pushed towards adipogenesis in chronic injury.

**Figure 5:**
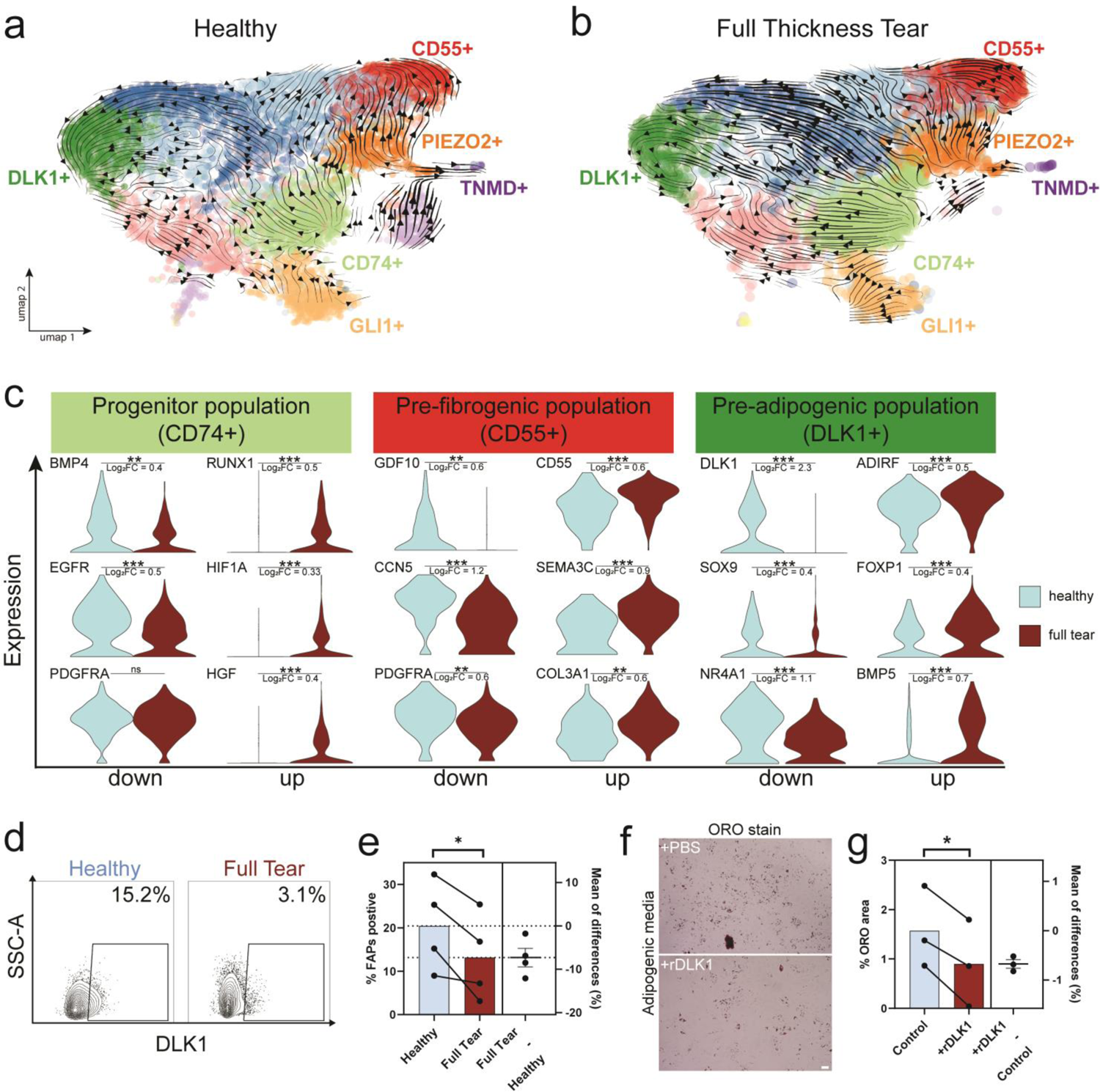
Chronic injury promotes proadipogenic gene and protein expression in FAPs. **a-b)** RNA velocity trajectory of FAPs derived from full thickness tear muscle samples (right panel) and corresponding healthy deltoid control muscle (left panel). UMAP is overlayed with arrows demonstrating predicted cell trajectories. Cells colored by FAP subpopulations. **C)** Violin plots depicting expression of select genes in the progenitor CD74+, pre-fibrogenic CD55+, and pre-adipogenic DLK1+ FAP subpopulations. Color indicates if FAPs derived from healthy (blue) or injured (red) muscle. Significant genes were Log_2_FC > 0.32 (fold change >1.25) and *p < 0.05 **p < 0.01 ***p < 0.001. **d)** Representative flow cytometry plots for DLK1 expression in FAPs from either healthy or full thickness tear muscle. **e)** Percentage of FAPs that are DLK1+ in healthy and full tear injured muscle. Paired patient samples are connected by lines. *p < 0.05. n = 4 biological replicates in d and e. **f)** Representative oil-red-o (ORO) images of FAPs grown in adipogenic media with PBS or recombinant DLK1 protein (rDLK1). Lipid is stained deep red. Scale bar = 40 µm. **g)** Quantification of the percentage of area stained for oil-red-o in FAP cultures grown in adipogenic media with PBS or rDLK1. Paired patient samples are connected by lines. n = 3 biological replicates in f and g. All plots are mean ± SEM. Statistical test for all plots: paired two-tailed t test. *p < 0.05. See also Supplementary Figure 6.

Given the pronounced trajectory changes, we sought to uncover the gene expression differences in the progenitor, pre-fibrogenic, and pre-adipogenic populations between healthy and full thickness injury muscle (**Figures 5c, S6b and Supplementary Table S7)**. Within the progenitor population, we found significantly increased expression of *RUNX1*, *HIF1A*, *FGF7*, and *HGF* in FAPs from full thickness injury muscle compared to healthy control muscle. The transcription factors RUNX1 and HIF1A have been shown to attenuate both fibroblast and adipocyte lineage specification^48–53^. FGF7 and HGF have known roles in regulating tissue regeneration via several downstream effects including activation of progenitor cells and immunomodulation^20,21,54–56^. These findings suggest that CD74+ cells upregulate genes that control stem/progenitor cell activation and lineage specification during injury. *BMP4* is down regulated in the injured CD74+ population compared to CD74+ cells from healthy muscle. BMP4 regulates adipogenesis by promoting white adipocyte browning^57,58^. Additionally, we found downregulation of *EGFR* in CD74+ cells. Low levels of EGF signaling induce pre-adipocyte differentiation in 3T3-L1 cells^59^. Together, this suggests that upon injury, progenitor cells activate differentiation programs and secrete factors that may drive FAP cell commitment to an adipogenic lineage.

In the pre-fibrogenic population, expression of genes associated with regulation of fibrosis, such as *CD55*, *SEMA3C*, and *CD70*, were increased in injured FAPs (**Figures 5c, S6b, and Supplementary Table S7**)^33,60–63^. *COL3A1*, which is important for fibrillogenesis, was also significantly increased^64,65^. The antifibrotic factors *GDF10* and *CCN5* were significantly decreased in injury^66–69^. GDF10 has also been demonstrated to be a negative regulator of adipogenesis in dystrophic muscles, which may act on FAP progenitor populations^70^. The extracellular matrix gene COL15A1 was also significantly decreased in injury. Interestingly, COL15A1 is decreased in fibrotic tissue from patients with Dupuytren’s contractures, another fibrotic musculoskeletal disease^71^. Furthermore, pre-fibrogenic FAPs from injured muscle have significantly less expression of *PDGFRα*, suggesting that these cells are pushed towards terminal differentiation. The altered expression of multiple secreted factors involved in fibrosis and adipogenesis in injury, suggests that CD55+ cells may be important in maintaining a healthy muscle environment. The increased proportion of pre-fibrogenic FAPs in chronic injury may lead to a loss of protective anti-fibrotic and anti-adipogenic signaling, which may ultimately lead to the promotion of fibrosis and adipogenesis.

Next, we hypothesized that the gene regulation within pre-adipogenic FAPs could further explain the increase in fatty infiltration seen in full thickness tear muscle. Akin to the pre-fibrogenic cells, injured pre-adipogenic FAPs had significantly less expression of *PDGFRα*, suggesting the FAPs differentiate and down regulate *PDGFRα*. Injured pre-adipogenic FAPs upregulated pro-adipogenic genes, including *ADIRF*, *FOXP1*, *BMP5*, and *TGFBI*, and downregulated negative regulators of adipogenesis, including *NR4A1*, *SOX9*, and *DLK1* (**Figures 5c, S6b and Supplementary Table S7**)^72–79^. IRF1 was also significantly decreased, which may further attenuate lineage specification away from fibroblasts^80,81^. IRF1 has been shown to promote fibrosis through downregulation of a key regulator of adipogenesis, C/EBPβ^80^, suggesting that the loss of IRF1 in injured pre-adipogenic FAPs would promote their differentiation towards adipogenesis rather than fibrosis^82^.

Remarkably, *DLK1* expression itself was robustly decreased in injured DLK1+ pre-adipogenic FAPs with an approximately 5-fold significant difference. A proposed mechanism for DLK1-mediated inhibition of adipogenesis is through upregulation of *SOX9* expression^27,45^. SOX9 inhibits adipogenesis through direct binding to the promoter regions of C/EBPβ and C/EBPδ^45^. We found that *SOX9* was down regulated in injured DLK1+ FAPs. Additionally, injured DLK1+ cells have higher levels of mature spliced *SOX9* mRNA than nascent mRNA, suggesting its transcription is decreased during injury (**Figure S6c**). Similarly, in injured FAPs, nascent *DLK1* mRNA is decreased along the trajectory from progenitor to pre-adipogenic cells (**Figure S6c**). This suggests that upon injury, downregulation of DLK1 leads to increased adipogenesis and fatty infiltration of skeletal muscle. Furthermore, the decrease in *HIF1A* expression in the CD74+ population may further attenuate DLK1-mediated repression of adipogenesis as HIF1A stabilizes the cleavage of membrane bound DLK1 leading to both extracellular and intracellular DLK1 signaling cascades (**Figure 5c**)^83^.

These results suggest that both intrinsic FAP subpopulation gene perturbations and FAP extrinsic changes in resident muscle cell gene signaling are driving fibro/fatty degeneration in chronic muscle injury. Taken together, the data from these experiments demonstrate a switch towards adipogenesis in injury conditions. Furthermore, in injury FAPs are pushed down the pre-adipocyte lineage but are likely transient as these cells become fat^29^. This in turn leads to overall less transient pre-adipogenic FAPs represented in injury as found in Figure 4. Injured FAP subpopulations upregulate genes that promote adipogenesis and downregulate genes that drive anti-adipogenic pathways.

### DLK1 prevents adipogenic differentiation of human fibroadipogenic progenitors

In our scRNA-seq data, we found expression of *DLK1* to mark a subset of CD10+ human pre-adipogenic FAPs. We confirmed that this transcriptionally-defined FAP subpopulation could be identified at the protein level, and found that there is a lineage trajectory of progenitor FAPs to DLK1+ pre-adipogenic FAPs. Furthermore, our data show that *DLK1* mRNA expression is significantly decreased in FAPs from chronically injured muscle. Therefore, we hypothesized that DLK1 protein is also decreased in chronic injury, leading to loss of DLK1-mediated inhibition of adipogenesis, which in part results in fatty degeneration of muscle. Given the pronounced decreased expression of *DLK1* mRNA in the pre-adipogenic population within the chronic full thickness tear injury muscle, we first sought to confirm that DLK1 protein expression was also decreased. We compared FAPs isolated from healthy deltoid muscle to FAPs from injured full thickness tears with full spectrum flow cytometry for DLK1 protein expression (**Figures 5d**). Akin to the scRNA-seq data, FAP specific DLK1 protein expression was significantly decreased in injured compared to healthy muscle (**Figure 5e**). We hypothesized that DLK1 loss results in increased adipogenic differentiation of FAPs and that this could be reversed through treatment with DLK1 protein. We tested this *in vitro* by culturing FAPs in adipogenic media for 1 week with recombinant DLK1 protein compared to PBS control (**Figure 5f**). Addition of DLK1 reduced adipogenic differentiation by greater than 50% (**Figure 5g**). Thus, DLK1 treatment results in FAP specific amelioration of adipogenesis in this *in vitro* model.

### DLK1 treatment inhibits fatty infiltration *in vivo*

We hypothesized that DLK1 treatment would also prevent adipogenesis and subsequently fatty infiltration of muscle after injury *in vivo*. Prior work has looked at the effect of DLK1 in mouse muscle injury with cardiotoxin and did not find a phenotype after DLK1 knockout^84^. However, this injury method does not produce an environment conducive to adipogenesis such as with glycerol injury^85^. In order to study the effect of DLK1 treatment on fatty infiltration after muscle injury we employed xenotransplantation of human FAPs along with glycerol injury (**Figure 6a**). Human FAPs suspended in 50% glycerol in sterile saline were transplanted into the tibialis anterior muscles of immunodeficient NSG mice. The mice were split into two groups which received either PBS or rDLK1 in the injection cocktail. The following day the same mice were injected with PBS or rDLK1 intramuscularly into the tibialis anterior muscles. The mice were analyzed after 14 days for the degree of fatty infiltration and the engraftment of human cells (**Figure 6b**). We quantified the amount of fat with immunofluorescence staining of perilipin, a marker of mature adipocytes. The mice treated with rDLK1 developed over 5-fold less fat that those receiving PBS (**Figure 6c**). In addition, we quantified the number of human cells within the mouse muscle through staining of lamin a/c, a human specific nuclear membrane marker. The average number of human cells engrafted in the rDLK1 group was over 3-fold more than the PBS group (**Figure 6d**). Taken together these data show that DLK1 treatment is sufficient to prevent fatty infiltration *in vivo* after injury. We also quantified myofiber cross section area and found that mice treated with rDLK1 had significantly less small rudimentary fibers suggesting reduced muscle degeneration in these mice (**Figure 6e and 6f**). We propose that loss of DLK1 protein expression in chronic full thickness rotator cuff injuries contributes to fatty degeneration in this patient population (**Figure 7**).

**Figure 6:**
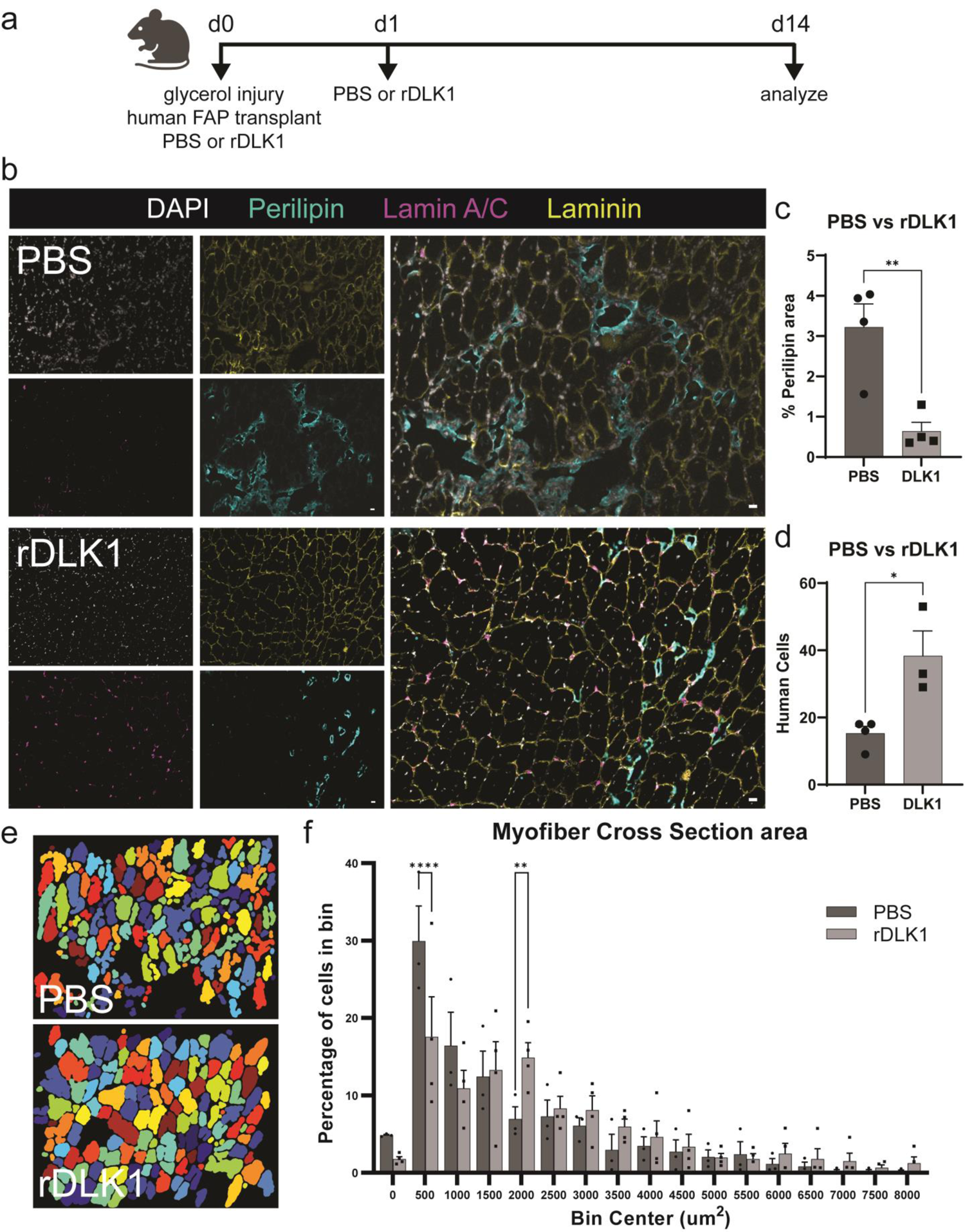
DLK1 treatment inhibits fatty infiltration *in vivo*. **a)** Schematic of experimental design for human FAP xenotransplantation and glycerol injury followed by rDLK1 treatment. At day 0 NSG mice were transplanted with human cells suspended in 50/50 glycerol in PBS with addition of either PBS or rDLK1. A second dose of intramuscular PBS or rDLK1 was administered on day 1. Mice were analyzed at day 14 post injury. **b)** Representative immunofluorescence cross section images of transplanted mouse tibialis anterior muscles treated with PBS or rDLK1, harvested at day 14 post injury. Nuclei marked with DAPI, adipocytes marked by perilipin, human nuclei marked by lamin a/c, and myofibers marked by laminin. Merged images on the right. Scale bars = 20 µm. **c)** Quantification of fatty infiltration by perilipin area in PBS vs. rDLK1 treated mice. **d)** Quantification of human cell engraftment by lamin a/c positive nuclei in PBS vs. rDLK1 treated mice. Statistical test for c-d: two-tailed t test. *p < 0.05 **p < 0.01. **e)** Masked images of myofiber cross section area utilizing laminin staining of mice treated with PBS vs. rDLK1. **f)** Quantification of binned myofiber cross section area of mice treated with PBS vs. rDLK1. Statistical test for e-f: two-tailed t test. **p < 0.01 ****p < 0.0001. All plots are mean ± SEM. n = 4 biological replicates (transplanted human FAPs derived from one human muscle sample) for all data.

**Figure 7:**
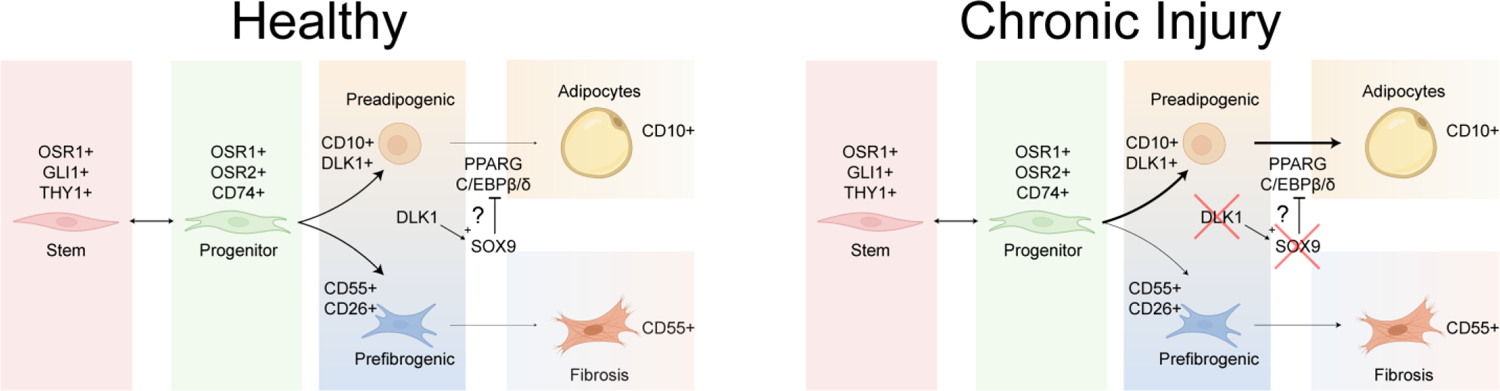
DLK1 prevents adipogenic differentiation of human fibroadipogenic progenitors. Model of proposed human FAP heterogeneity and lineages in healthy muscle and during chronic injury. CD55 is increased along fibrogenic differentiation while CD10 is increased along adipogenic differentiation. DLK1 both marks preadipogenic FAPs and prevents adipogenic differentiation possibly through increased expression of SOX9.

## Discussion

In this study, we sought to investigate stem cell functional heterogeneity in a clinically relevant model of human muscle disease. Our dataset provides a valuable resource of over 24,000 fibroadipogenic progenitor cells, and over 14,000 cells of other major resident cell types of healthy and injured skeletal muscle. We utilized scRNA-seq to identify human FAP subpopulations and propose that these are functionally heterogeneous. We highlight five clusters which likely serve functionally distinct roles in homeostasis and regeneration: progenitor (GLI1+ and CD74+), pre-adipogenic (DLK1+), pre-fibrogenic (CD55+), and transitional mechanosensitive (PIEZO2+) subpopulations. Utilizing full spectrum flow cytometry, we were able to resolve these unique stem and progenitor cell populations through surface markers and provide a roadmap for prospective isolation as full spectrum flow sorting machines become more widely available^86^. We then show that FAP subpopulations are perturbed in injury and these changes are highly sensitive to injury severity. In severe chronic injury, subpopulation gene expression and cell proportions are both affected with the net result of increased FAP-derived adipocytes, mirroring the clinical phenotype. Finally, we propose that DLK1 is both a marker of pre-adipogenic cells and is a crucial regulator of FAP adipogenesis. DLK1 protein is sufficient to significantly inhibit FAP commitment to adipogenic differentiation, possibly through upregulation of SOX9. Furthermore, treatment of DLK1 *in vivo* prevents fatty infiltration of muscle after injury and reduced myofiber degeneration.

OSR1 is thought to mark true FAP progenitor populations in mice^17,19,38^. We identified two distinct human FAP OSR1+ populations which express high amounts of either *GLI1* or *CD74*. The GLI1+ population expressed high levels of hedgehog signaling markers and *THY1*. Both *GLI1* and *THY1* expressing cells are highly proliferative and responsible for *de novo* adipogenesis in skeletal muscle^9,85^. The supply of hedgehog ligand has yet to be confirmed in humans but is thought to come from Schwann or endothelial cells^85,87^. Interestingly we found high expression of the hedgehog signaling protein desert hedgehog (*DHH*) in a subset of endothelial cells which may be the cellular origin of this ligand in human skeletal muscle (**Figure S7a**). In the intestine, PDGFRα+ subepithelial mesenchymal cells that express GLI1 maintain the epithelial stem cell population through secretion of Wnt2b or Wnt3a^88^. The human GLI1+ FAP population identified here also has specific and high expression of *WNT2b* (**Figure S7b**). FAPs may regulate the stem cell niche through the secretion of WNT ligands from a specific hedgehog-responsive FAP subpopulation.

In muscle injuries, progenitor cells give rise to new populations that prevent proper regeneration^4,9^. Treatments that can prevent progenitors from giving rise to degenerative cell populations could improve patient outcomes after injury. For example, treatment of FAPs with a beta-agonist induces FAP beigeing, a process thought to produce specialized adipocytes that are regenerative and thus improves muscle regeneration after injury^39^. *CXCL12* was upregulated in beigeing FAPs and is also a marker of activated CD74+ FAPs in our dataset (**Figure S7c**). We hypothesize that CXCL12 is critical for directing FAP progenitor fate and may function through CXCL12 – CXCR4 signaling, as expression of *CXCR4* is present on both muscle stem cell and immune cell populations (**Figures S7a**)^89,90^. By uncovering novel markers of FAP subpopulations, we can begin to unravel the mechanisms by which either pro-regenerative or degenerative FAP states are produced. It will be critical for future studies to investigate the “tunability” of this population to influence FAP differentiation towards pro-regenerative subpopulations and away from degenerative adipogenic and fibrogenic states.

We further demonstrate that utilization of full spectrum flow cytometry enables highly specific characterization of stem cell populations in skeletal muscle. Identification of surface markers of pre-fibrogenic and pre-adipogenic FAP populations provides a method for the direct study of FAP heterogeneity both *in vivo* and *in vitro*. CD55 marks fibrogenic FAPs; while, DLK1 and CD10 mark pre-adipogenic FAPs and CD10 expression is increased along adipogenic differentiation. Interestingly, DLK1 expression was decreased *in vitro*. DLK1 expression is decreased by dexamethasone, a common component in adipogenesis treatment media^28^. We propose that DLK1 marks pre-adipogenic FAPs but is lost upon adipogenic differentiation. The identification of these surface markers offers an attractive way to prospectively study and isolate subpopulations including fibrogenic and adipogenic FAPs.

We next applied this knowledge of FAP subpopulations to a chronic injury model of rotator cuff injuries. We found underlying gene and protein expression was altered depending on whether the FAPs were isolated from muscles with partial or full tendon injuries, suggesting that FAPs respond uniquely to the severity of the rotator cuff tendon tear. As injury severity increases (partial to full tear) FAP lineage specification switches to a predominantly adipogenic cellular program. This correlates to the clinical phenotype, with increased severe fatty infiltration on MRIs of patients with larger, chronic rotator cuff tears^40^. Indeed, there appears to be FAP subpopulation specific changes that while unique to each population, are all leading to a net increase of adipogenesis in response to injury severity.

There are currently no accepted pharmacologic strategies to prevent muscle degeneration. FAP subpopulation-specific changes in injury offer novel avenues for therapeutics development. For example, in rotator cuff injuries the degree of muscular fatty degeneration is negatively correlated with clinical outcomes even after successful tendon repair. When degeneration becomes severe, repair is overwhelmingly likely to fail, thus these patients must undergo a more extreme treatment consisting of reverse total shoulder replacement to restore shoulder function^42,91,92^. Even mild levels of fatty degeneration persist after repair, highlighting the need for adjunctive medical treatments^92^. As such, intervening before degeneration becomes advanced is of paramount importance. We propose the CD74+/OSR1+/OSR2+ population is the human progenitor population that can be targeted in injury to ameliorate fatty degeneration. Even in chronic tears, there appear to be an abundant number of these cells in supply, despite the overall switch to a pro-adipogenic program, which could be reprogrammed to promote muscle regeneration.

We see more evidence for a pro-adipogenic program switch within the pre-adipogenic FAP population. We found DLK1 expression is greatly decreased in pre-adipogenic FAPs in injured muscle. One proposed mechanism for the anti-adipogenic effects of DLK1 is through upregulation of *SOX9*, which was significantly decreased in injured cells within this population^27,45^. Moreover, we were able to confirm that DLK1 protein expression is decreased in FAPs from chronic full thickness tears. The loss of DLK1 expressing cells could at least in part explain the fatty infiltration seen in the muscle from these patients, through either commitment and differentiation of these cells towards adipocytes or loss of DLK1-mediated suppression of adipogenesis. Indeed, DLK1 treatment of human FAPs can inhibit FAP adipogenesis even in pro-adipogenic environments both *in vitro* and *in vivo*. Our xenotransplant experiments are the first to show that human FAPs can be engrafted into an immunodeficient mouse muscle. Engraftment of human FAPs is increased and fatty infiltration is reduced by rDLK1 treatment after muscle injury, demonstrating the capability of DLK1 to suppress adipogenesis.

Thus, we propose a model of muscle degeneration where DLK1 is a part of a negative feedback loop to prevent commitment of pre-adipogenic FAPs to adipogenesis. These findings demonstrate the importance of understanding stem and progenitor cell heterogeneity within musculoskeletal injuries. This dataset can provide a valuable resource for the identification of molecular targets to direct progenitors towards pro-regenerative fates and suppress the production of degenerative phenotypes. It will be critical for future translative studies to utilize this knowledge of human FAP heterogeneity to target specific cell states and populations in degenerative diseases.

## Acknowledgments

We thank Dr. Lauren E. Byrnes for her thoughtful review and comments, the UCSF Parnassus Flow CoLab for assistance with flow cytometry experiments (RRID:SCR_018206 and supported in part by Grant NIH P30 DK063720 and by the NIH S10 Instrumentation Grants S10 1S10OD026940-01 and S10 1S10OD021822-01), the UCSF Genomics CoLab for assistance with scRNA-seq experiments, the UCSF Wynton high performance computing cluster for analysis, and the following grants: OREF A137519 to S.M.G., NIH 3R01AR072669 and VA grant 1 I01 CX002200-01A2 to X.L. and B.T.F, NIH 5R01AR072669-04 to S.M.G and B.T.F., and NIH 3R01AR072669 - 05S2 to S.M.G and B.T.F. Figures 1a, 2b/e/j, 3k, and 7 made with BioRender.

## Author Contributions

Conceptualization, S.M.G. and B.T.F.; Methodology, S.M.G. and B.T.F.; Software, S.M.G. and J.L.; Validation, S.M.G. and B.T.F.; Formal Analysis, S.M.G., J.L., and A.D.; Investigation, S.M.G., J.L., A.D., H.C., and M.L.; Data Curation, S.M.G.; Writing – Original Draft, S.M.G.; Writing – Review & Editing, S.M.G., J.L., A.D., X.L., M.R.D., and B.T.F.; Visualization, S.M.G. and J.L.; Supervision, X.L. and B.T.F.; Project Administration, X.L. and B.T.F.; Funding Acquisition, S.M.G., X.L., and B.T.F.

## Inclusion and diversity

We support inclusive, diverse, and equitable conduct of research.

## Materials and Methods

### Muscle Procurement and Digestion

This study was conducted under the approval of the Institutional Review Board (IRB # 18-26000 and 18-26760) at The University of California San Francisco (UCSF). Biopsies were obtained from individuals undergoing surgery at UCSF. Written informed consent was obtained from all subjects. Deltoid and supraspinatus muscle samples were obtained during arthroscopic surgery with a biting grasper tool. Hamstring muscle was obtained from harvested hamstring tendons utilized for anterior cruciate ligament reconstruction. All samples were transferred from the operating room on ice and stored in 30% FBS (fetal bovine serum, Gibco, #10437028) in Dulbecco’s Modified Eagle Medium (DMEM) (Gibco, #11965084) until processing. Muscle was trimmed of excess fat, tendon, and connective tissue. Muscle was then mechanically and enzymatically digested in 1 mg/ml collagenase III (Worthington Biochemical Corporation, #NC0021929) in DMEM with 10% FBS and 1% Penicillin/Streptomycin (Gibco, #15140122) at 37°C for 70 minutes with intermittent manual needle trituration with an 18-gauge needle^93^. Digests were washed with PBS (Gibco, #10010023) and further digested with 0.25% trypsin-EDTA (Gibco, #25200056) at 37°C for 12–15 min. Suspensions were passed through 40 µm nylon mesh (Corning, #431750), erythrocytes were lysed with ACK lysing buffer (Quality Biological, #188-156-101) for 5–7 min on ice and washed in PBS with 2% FBS. Cells were washed and resuspended in 100 µl flow cytometry buffer (PBS with 2% FBS, 5 mM EDTA (Corning, #MT-46034CI), 2 mM HEPES (Quality Biological, #118-089-721)) per 1 gram of starting tissue weight. Antibody cocktail consisting of: anti-CD31 (eBioscience, #48-0319-42, eFluor 450) (2 µl per gram of starting muscle weight), anti-CD45 (eBioscience, #48-0459-42, eFluor 450) (2 µl), anti-CD56 (Miltenyi Biotec, #130-114-548, APC-Vio770) (8 µl), anti-CD34 (BD, #555823, PE-Cy5) (6 µl) and anti-PDGFRα (BioLegend, 323506, PE) (8 µl) was then added to the cells for 30 minutes on ice. Cells were washed and resuspended in flow cytometry buffer with 1:1000 SYTOX Blue (ThermoFisher, #S34857). Flow cytometry analysis and cell sorting were performed at the UCSF Flow Cytometry Core with BD FACSAria2 machines operated using FACSDiva software. Flow cytometry isolations were analyzed with FACSDiva and FlowJo software.

### Single Cell mRNA Sequencing

Viable CD31-/CD45-/CD56-/CD34+/PDGFRa+ FAPs were collected and spiked into a collection of all live (SYTOX-)cells to obtain samples with >50% FAPs. Cells were resuspended in 0.04% BSA (EMD Millipore, #12659-500gm) in PBS at 1000 cells per μl to prepare for scRNA-seq. For each sample 5,000-10,000 cells were loaded onto one lane of a 10X Chromium chip per sample, using the Chromium Single Cell 3’ Reagent kit (10X Genomics, version 3.1, #PN-1000123). Manufacturer’s instructions were followed for Gel Beads-in-emulsion (GEM) generation, cDNA production and library construction. Libraries were sequenced on Illumina NovaSeq. CellRanger (version 3.0) was used with default settings to de-multiplex, align reads to the human genome (10X Genomics pre-built GRCh38 reference genome) and count unique molecular identifiers (UMIs). Doublets were removed from individual datasets with DoubletFinder based on loading concentrations and the 10X Genomics estimates for doublet rates^94^. Seurat (version 4.1) was used for downstream analysis of individual datasets, including filtering, dimension reduction, clustering, UMAP, and differential gene expression analysis (see GitHub repository for code)^95^. Samples were integrated according to the Seurat Integration Tutorial. Scaling, dimension reduction, and clustering were performed in Seurat to integrate data for visualization. Differential gene expression analysis and gene expression visualization were performed in Seurat on the original, non-integrated values. For pseudotime analyses, UMAP dimensions from Seurat were imported into Monocle3 (version 1.0)^96^. Pseudotime ordering and branch analysis was then performed according to the Monocle3 tutorial. RNA velocity analyses performed with scVelo according to the Differential Kinetics tutorial (version 0.2.5)^47,97^.

### Full spectrum flow cytometry

For analysis of freshly harvested samples, muscle was digested, and cell suspensions were prepared as above. Cells were stained with anti-CD31 (eBioscience, #63-0319-42, Super Bright 600) (2 µl per gram of starting muscle weight), anti-CD45 (eBioscience, #48-0459-42, eFluor 450) (2 µl), anti-CD56 (Miltenyi Biotec, #130-114-548, APC-Vio770) (8 µl), anti-CD34 (BD, #555823, PE-Cy5) (6 µl), anti-THY1 (eBioscience, #11-0909-42, FITC) (2 µl), anti-CD74 (BioLegend, #326808, PE) (2 µl), anti-DLK1 (R&D, #FAB1144N, Alexa Fluor 700) (4 µl), anti-CD10 (BioLegend, #312226, BV711) (2 µl), anti-CD26 (BioLegend, #302710, APC) (1 µl), and anti-CD55 (BioLegend, #311314, PE-Cy7) (2 µl). We utilized single color controls both with both beads (ThermoFisher, #01-1111-42) and cells. For analysis of cultured FAPs, cells were gently lifted with 0.05% trypsin, washed, resuspended in 100 µl flow cytometry buffer (PBS with 2% FBS, 5 mM EDTA, 2 mM HEPES) per 1×10^6^ cells, and stained with anti-THY1 (2 µl per 1×10^6^ cells), anti-CD74 (2 µl), anti-DLK1 (4 µl), anti-CD10 (2 µl), anti-CD26 (1 µl), and anti-CD55 (2 µl). Fluorescence minus one (FMO) controls were made with one marker left out of the antibody cocktail for each respectively. In both fresh digests and cultured cell preparations, cells were washed and resuspended in flow cytometry buffer with 1:2000 SYTOX Blue viability dye (ThermoFisher, #S34857) added just prior to analysis. Flow cytometry analysis and cell sorting were performed at the UCSF Flow Cytometry Core with Cytek Aurora machines operated using SpectroFlo software. Flow cytometry data were analyzed with SpectroFlo and FlowJo software. Dimension reduction was performed in FlowJo with UMAP^37^. Data was then visualized in R with ggplot2.

### FAP culture

FAPs were isolated from fresh tissue digests with the surface marker combination CD31-/CD45-/CD56-/CD34+. Primary cells were cultured in expansion media: Ham’s F-10 Nutrient Mix (Gibco, #11550043) containing 1% Penicillin/Streptomycin, 10% FBS, and 10ng/mL bFGF (Gibco, #PHG0264). Cell media was changed every 2-3 days and cells were passaged at 70-80% confluency. For adipogenic differentiation, FAPs were grown to 80% confluency in expansion media and then switched to adipogenic media (StemPro Adipogenesis Differentiation Kit, Gibco, # A1007001). Media was changed every 2 days. For fibrogenic differentiation, FAPs were grown to 80% confluency in expansion media and then switched to expansion media containing, 10 ng/ml of TGFβ1 (Invitrogen, #PHG9204). Media was changed every 2 days.

### Immunofluorescence and imaging

Human muscle biopsies were obtained from the operating room and transferred on ice. Tissues were flash frozen within O.C.T. Compound (Tissue-Tek, #4583) in liquid nitrogen cooled 2-methylbutane (Millipore Sigma, #MX0760) and stored at −80 degrees. Transverse muscle sections were cut at 8 µm and stored at −80 degrees. Slides were warmed at room temperature prior to initiating staining protocols. Slides were fixed in 4% PFA (Electron Microscopy Sciences, #50-980-487) at room temperature for 10 min, washed in PBST (PBS with 0.1% Tween-20 (Millipore Sigma, #P1379)), then blocked with protein-free serum block (Agilent, #X090930-2) for 1 hour and incubated at 4 degrees overnight with the corresponding primary antibodies. Primary antibodies and dilutions for human muscle cross section imaging: anti-human PDGFR alpha antibody 1:100 (R&D Systems, #AF-307-NA), anti-Laminin 2 alpha 1:250 (Abcam, #ab11576), anti-CD55 1:100 (Invitrogen, #MA1-911610), anti-CD90 (Thy-1) 1:100 (Invitrogen, #14-0909-82), and anti-DLK-1 1:100 (Abcam, #ab89908). After a PBST wash the corresponding secondary antibodies were applied for 1 hr at room temperature. Secondary antibodies and dilutions for human muscle cross section imaging: Donkey anti-mouse (Alexa Fluor® 488) 1:500 (Abcam, #ab150105), donkey anti-goat (Alexa Fluor® 594) (Invitrogen, #A-11058) 1:500, and donkey anti-rat (Alexa Fluor® 647) 1:500 (Abcam, #ab150155). After a PBST wash, slides were mounted with VECTASHIELD mounting medium with DAPI (Vector Laboratories, #H-2000-2). For immunofluorescence imaging of cultured cells, cells were washed with PBS and fixed with 4% PFA at room temperature for 10 min, and then washed, blocked, and stained in the same manner as with slides described above. Primary antibodies and dilutions used for stains of cultured cells: anti-CD10 1:100 (Santa Cruz Biotechnology, INC., #sc-19993) and anti-CD55 1:100 (Invitrogen, #MA1-911610). Secondary antibodies and dilutions used for stains of cultured cells: Donkey anti-mouse (Alexa Fluor® 488) 1:500 (Abcam, #ab150105). Cells were counter-stained with DAPI 1:10,000 (Millipore Sigma, #D9542-10mg) for 5 minutes at room temperature, then washed with PBST, and kept in PBS for imaging. For quantification of lipid production in cell cultures, cells were fixed with 4% PFA at room temperature for 10 minutes, rinsed three times for 30 seconds in DI water, and then stained with 0.36% Oil Red O solution (EMD Millipore, #90358) for 30 minutes. Cells were once again rinsed three times for 30 seconds in DI water and then kept in 10% glycerol in PBS for imaging. All images were taken on a Zeiss Axio Observer D1 fluorescence microscope and were analyzed using ImageJ.

### *In vitro* DLK1 treatment

FAPs were seeded on polystyrene plates at a density of 60,000 cells/cm^2^. At 80% confluence cells were treated with adipogenic media (Gibco, #A1007001) containing 5 µg/ml rDLK1 resuspended in PBS (R&D Systems, #1144-PR-025) or PBS vehicle. Media was changed every 2 days.

### Animal housing and Xenotransplant glycerol-induced injury model

All animal procedures were approved by the San Francisco VA Medical Center Institutional Animal Care and Use Committee and performed under IRB protocol. All experiments were carried out on three-month old male immunodeficient NOD.Cg-Prkdcscid Il2rgtm1Wjl/SzJ from The Jackson Laboratory. Mice were maintained on a standard chow diet with 12-hour light and dark cycles. Mouse tibialis anterior muscles were injected with 50 uL solution of 50,000 human FAPs resuspended in 50% glycerol in PBS with either 29ng/ul concentration of recombinant DLK1 resuspended in PBS (R&D Systems, #1144-PR-025) or with control (identical volume of PBS) via insulin syringe. Injections of PBS or rDLK1 were repeated 24 hours later. The tibialis muscles were then harvested after 14 days.

### Mouse Tibialis anterior immunofluorescence staining and analysis

Tibialis anterior muscles were flash-frozen in optimum cutting temperature compound (OCT) in liquid nitrogen-cooled isopentane and sectioned (8um) with a cryostat in cross section orientation serially through the entire muscle. Slides were fixed in 4% paraformaldehyde for 10 minutes at room temperature, washed with 0.1% PBST (Phosphate Buffered Saline Tween 0.1%), and blocked in 5% bovine serum albumin for 30 minutes. Samples were then incubated with laminin 2 alpha 1:500 (Abcam, #ab11576), lamin-AC 1:100 (Abcam, #MA3-1000), Perilipin A 1:250 (Sigma-Aldrich, #P1998) at 4°C overnight. Slides were then rinsed and incubated with secondary antibodies at 1:250: donkey anti-rabbit IgG Alexa Fluor 488 (Invitrogen, #A-21206), donkey anti-mouse Alexa Fluor 594 (Invitrogen, #A-21203), donkey anti-rat Alexa Fluor 647 (Abcam, #ab150155) at 1 hour at room temperature. After a PBST wash, slides were mounted with VECTASHIELD mounting medium with DAPI (Vector Laboratories, #H-2000-2). All images were taken on a Zeiss Axio Observer D1 fluorescence microscope and were analyzed using ImageJ. Myofiber cross section area analysis was performed in CellProfiler utilizing the Muscle2View pipeline^98^.

### MRI imaging and analysis

T2-relaxtion magnetic resonance imaging studies were utilized to grade muscle fatty infiltration in the supraspinatus and deltoid muscles as originally described by Goutallier et al^41^. Classification as follows: Goutallier 0: no appreciable fat content, Goutallier 1: some fat streaking, Goutallier 2: fat is appreciable but less fat than muscle, Goutallier 3: fat content equal to muscle, Goutallier 4: fat content is greater than muscle.

### Statistical Testing

All statistical testing was performed using Prism 9.4 software. Statistical tests and p-value cut-offs used for each experiment are listed under the relevant heading.

### Resource Availability Lead Contact

Further information and requests for resources and reagents should be directed to and will be fulfilled by the lead contact, Brian Feeley.

## Materials Availability

This study did not generate new unique reagents.

## Data and Code Availability

Raw and processed scRNA-seq data available on NIH GEO (https://www.ncbi.nlm.nih.gov/geo/) series number to follow publication. scRNA-seq data will be deposited on CellxGene. Code and scripts utilized in this study are available at: https://github.com/StevenMGarcia/Feeley-Liu_Lab

## Supplemental Figure

Figure S1: Supplementary data for figure 1. Single cell sequencing identifies human fibroadipogenic progenitor populations. **a)** Representative flow cytometry plots of human FAP sorting scheme. FAPs are CD45-/CD31-/CD56-/CD34+/ PDGFRα+. CD45-/CD31-/CD56-/CD34+ cells are >95% PDGFRα+. Inset values are percentage of cells in each gate respectively. **b)** UMAP of all cells obtained from 14 patient samples, separated by patient and sample, and labeled by age and sex in black text. Total cell numbers for each sample in black text. Cells are colored by cell type and number of cells in each cluster labeled in the same color. **c)** Expression of localized gene expression of cell type specific markers for FAPs (*PDGFRA*), immune cells (*PTPRC* also known as *CD45*), macrophages (*CD68*), T-cells (*CD3E*), endothelial cells (*PECAM1* also known as *CD31*), pericytes (*RGS5*), myofibroblasts (*ACTA2*), and myogenic cells (*PAX7* and *MYOD1*). **d)** UMAP of FAPs obtained from 14 patient samples, separated by patient and sample, and labeled by age and sex in black text. Total cell numbers for each sample in black text. Cells are colored by cell type and number of cells in each cluster labeled in the same color. **e)** Gene expression within FAP subpopulations. **f)** Expression of each of the markers shown in e.

Figure S2: Supplementary data for figure 1. Distinct human fibroadipogenic progenitor populations are also marked by *CCN3*, *LAMA2*, *ATF3*, *MYL9*, *TNMD*, and *TNNT1*. a) Model UMAP of FAP subpopulations with highlighted populations. b) Expression of gene markers the highlighted populations. Deeper red color indicates higher expression.

Figure S3: Supplementary data for figure 2. Transcriptionally heterogenous human FAPs also have heterogenous protein expression of distinct surface markers. **a)** Representative full spectrum flow cytometry analysis of human FAPs. The antibody cocktail included antibodies against CD31, CD45, CD56, CD34, PDGFRα, THY1, CD74, DLK1, CD10, CD55, CD26, and a SYTOX live dead stain. **b)** Representative flow cytometry plots of fluorescence minus one (FMO) control samples for full spectrum flow cytometry analyses shown in figure 2c. Human FAPs identified by the marker combination CD45-/CD31-/CD56-/CD34+, stained with the full antibody cocktail minus one antibody respectively. **c)** Representative UMAP demonstrating overlap of CD26 and CD55 expression as well as overlap of DLK1 and CD10 expression. Cells gated as positive are colored in red. **d)** Total expression of each surface marker on FAPs utilizing CD10 (MME) in place of DLK1 as compared to figure 2g. DLK1 is more selective than CD10. **e)** Percentage of FAPs with each surface marker combination utilizing CD10 instead of DLK1 as one of the 4 markers, showing all possible combinations. **f)** Percentage of FAPs with each surface marker combination utilizing 4 markers showing all possible combinations of the data shown in figure 2g. n = 6 biological replicates for all data. All plots represent mean ± SEM.

Figure S4: Supplementary data for figure 3. DLK1/CD10 and CD55 are upregulated during FAP differentiation towards adipocyte or fibroblast cell fates. a) Pseudotime analysis demonstrating pseudotime values for cells colored on the UMAP. **b)** Expression along pseudotime for each lineage from progenitor cells down adipogenic, fibrogenic, and tenogenic pathways. Cells placed along pseudotime for each path depicted above the plots. Pseudotime split into 20 equal sized bins for each path. **c)** UMAP of scRNA-seq of human FAPs cultured for 3 weeks in three conditions: standard expansion media (std), adipogenic media (adipo), and fibrogenic media (fibro). **d)** Expression of markers of FAPs (*PDGFRα*), adipocytes (*PLIN1*), and fibroblasts (*ACTA2*). **e)** Violin plots of gene expression within cells of each type separated by culture condition. **f)** Violin plots of CD10 and CD55 expression within *PDGFRA* positive cells in each media condition. **g)** Total surface marker expression on FAPs cultured in each media compared to freshly harvested FAPs. **h)** UMAP of DLK1/CD10 double positive cells. **i)** Cells expressing all combinations of CD10 and CD55, compared to freshly harvested FAPs. n = 6 biological replicates for all data. All plots represent mean ± SEM. Statistical test for all plots: two-way ANOVA with Tukey’s multiple comparison correction. *p < 0.05 **p < 0.01 ***p < 0.001 ****p < 0.0001.

Figure S5: Supplementary data for figure 4. Severe chronic injury increases adipogenic and fibrotic commitment of FAPs. a) Summary of all patients’ supraspinatus and deltoid muscle fibro/fatty infiltration utilizing the Goutallier classification. **b)** UMAP of cells from healthy deltoid tissue compared to injured supraspinatus muscle. Cell number for each cluster in text colored by cluster. **c-d)** The proportion of each cluster in paired healthy and injured muscle samples for: **c)** partial thickness tendon tears and **d)** full thickness tendon tears compared to healthy deltoid muscle from the same patients. Color indicates if cells are from healthy (blue) or injured (pink) muscle sample. Paired samples linked with lines. n = 3 partial and n = 3 full thickness tear biological replicates. Statistically significant tested with a paired two-way ANOVA. *p < 0.05 **p < 0.01 ***p < 0.001 ****p < 0.0001.

Figure S6: Supplementary data for figure 5. FAP subpopulation gene expression in chronic injury leads to a predominantly proadipogenic tissue state. **a)** RNA velocity trajectory of partial thickness tear muscle. FAP subset UMAP is overlayed with arrows demonstrating predicted cell trajectories. Cells colored by FAP subpopulations. **b)** Violin plots of gene expression in the progenitor CD74+, pre-fibrogenic CD55+, and pre-adipogenic DLK1+ populations. Significant genes were Log_2_FC > 0.32 and *p < 0.05 **p < 0.01 ***p < 0.001. **c)** RNA velocity plots of healthy and full thickness tear muscle samples. Left boxed plots: plots of the spliced to unspliced RNA content in FAPs with learned dynamics overlayed. Right plots: velocity plots of the rate of change of unspliced to spliced RNA projected onto the FAP UMAP. Green = high rate of change, red = low rate of change.

Figure S7: FAPs promote muscle regeneration or fibro/fatty degeneration by expression and secretion of ligands that interact with neighboring FAPs and muscle cells. a) Gene expression by muscle cell type. b) WNT signaling ligand expression by FAP subpopulation. c) CXCL12 expression by FAP subpopulations. In all plots larger circle and darker red coloring depict larger proportion of cells expressing each corresponding gene and higher expression respectively.

## Supplemental Table

Supplemental Table S1: Related to Figure 1. Differential gene expression of integrated datasets including all cell types.

Supplemental Table S2: Related to Figure 1 and 2. Differential gene expression of human FAP subpopulations.

Supplemental Table S3: Related to Figure S4. Differential gene expression of cultured human FAPs by media condition.

Supplemental Table S4: Related to Figure S4. Differential gene expression of cultured human FAPs by cell type.

Supplemental Table S5: Related to Figure 5. Differential gene expression of FAPs from healthy vs. partial thickness tear injury muscle. Upregulated genes in healthy are > 0 and upregulated genes in injury are < 0.

Supplemental Table S6: Related to Figure 5. Differential gene expression of FAPs from healthy vs. full thickness tear injury muscle. Upregulated genes in healthy are > 0 and upregulated genes in injury are < 0.

Supplemental Table S7: Related to Figure 6. Differential gene expression of FAP subpopulations from healthy vs. full thickness tear injury muscle. FAP subpopulations separated by tabs. Upregulated genes in healthy are > 0 and upregulated genes in injury are < 0.

